# Multiplexed imaging of high density libraries of RNAs with MERFISH and expansion microscopy

**DOI:** 10.1101/238899

**Authors:** Guiping Wang, Jeffrey R. Moffitt, Xiaowei Zhuang

## Abstract

As an image-based single-cell transcriptomics approach, multiplexed error-robust fluorescence *in situ* hybridization (MERFISH) allows hundreds to thousands of RNA species to be identified, counted and localized in individual cells while preserving the native spatial context of RNAs. In MERFISH, RNAs are identified via a combinatorial labeling approach that encodes RNA species with error-robust barcodes followed by sequential rounds of single-molecule FISH (smFISH) to read out these barcodes. The accuracy of RNA identification relies on spatially separated signals from individual RNA molecules, which limits the density of RNAs that can be measured and makes the multiplexed imaging of a large number of high-abundance RNAs challenging. Here we report an approach that combines MERFISH and expansion microscopy to substantially increase the total density of RNAs that can be measured. Using this approach, we demonstrate accurate identification and counting of RNAs, with a near 100% detection efficiency, in a ~130-RNA library composed of many high-abundance RNAs, the total density of which is more than 10 fold higher than previously reported. In parallel, we demonstrate the combination of MERFISH with immunofluorescence. These advances increase the versatility of MERFISH and will facilitate its application of a wide range of biological problems.

## Introduction

*In situ* imaging-based approaches to single-cell transcriptomics allow not only the expression profile of individual cells to be determined but also the spatial positions of individual RNA molecules to be localized. These approaches provide a powerful means to map the spatial organizations of RNAs inside cells and the transcriptionally distinct cells in tissues^1^. Multiplexed fluorescence *in situ* hybridization (FISH)^2–7^ and *in situ* sequencing^8,9^ have been used to profile the expressions of RNAs in single cells. Among these, MERFISH, a massively multiplexed form of smFISH, enables RNA imaging at the transcriptomic scale with high accuracy and detection efficiency^7^. By imaging single RNA molecules, smFISH provides the precise copy number and spatial distribution of individual RNA species in single cells^10,11^. MERFISH multiplexes smFISH measurements by labeling RNAs combinatorically with oligonucleotide probes which contain error-robust barcodes and then measuring these barcodes through sequential rounds of smFISH imaging. Using this approach, we demonstrated simultaneous imaging of hundreds to thousands of RNA species in individual cells using barcoding schemes capable of detecting and/or correcting errors^7^. Recently, we have increased the measurement throughput of MERFISH to tens of thousands of cells per single-day-long measurement^12^. In addition, we developed a matrix-imprinting-based sample clearing approach that increases the signal-to-background ratio by anchoring RNA molecules to a polymer matrix and removing other cellular components that give rise to fluorescence background^13^. This clearing approach enabled high-quality MERFISH measurement of tissue samples^13^.

In order to accurately identify RNA molecules, MERFISH, as well as other multiplexed smFISH-based RNA profiling methods, requires non-overlapping signals from individual RNAs. However, when two molecules are sufficiently close to each other, the localization obtained from sequential rounds of smFISH imaging for one molecule will overlap with that from the other molecule, diminishing our ability to identify these RNAs and, thus, limiting the density of RNAs that can be profiled. This problem could potentially be mitigated by super-resolution optical imaging^14,15^, by analysis methods to address partially overlapping fluorophores^16–19^, or by sample expansion^20,21^. In particular, since neighboring RNA molecules may physically overlap in space, expansion microscopy (ExM), which uses sample expansion to effectively increase the distances between neighboring molecules^20^, may provide an especially effective means to increase the RNA density limit accessible by MERFISH. In ExM, the desired signal is conjugated to an expandable polyelectrolyte gel, and then the gel is physically expanded by changing the ionic strength of the buffer^20^. ExM has recently been combined with smFISH to help better resolve highly expressed RNAs, with either single-round or multiple rounds of smFISH to measure one or several genes^21,22^. In addition, RNAs have been anchored to a non-expandable polyacrylamide matrix to facilitate sample clearing and background removal in RNA FISH^13,23,24^ and MERFISH measurements^13^. Thus, we reason that ExM should also be compatible with MERFISH and may help substantially increase the RNA density measurable by MERFISH.

In this paper, we report an approach to combine MERFISH and ExM. We anchor mRNAs to an expandable polyelectrolyte gel via acrydite-modified poly-dT locked nucleic acid oligonucleotides hybridized to the polyA tail of mRNAs. We demonstrate the efficacy of this approach by imaging a high-abundance RNA library, which contains ~130 RNA species with a total RNA abundance that is 14-fold higher than what has been previously demonstrated with MERFISH imaging^12,13^, in cultured human osteosarcoma (U-2 OS) cells. Unlike our previous MERFISH measurements of lower-density RNA libraries, in which we demonstrated 80-90% detection efficiency, the RNAs in this high-density library are not well resolved and hence are detected with a relatively low detection efficiency of ~20% without gel expansion. In contrast, individual RNA molecules become well resolved in expanded samples, leading to a substantial increase in their detection efficiency. Comparison with smFISH and bulk sequencing results demonstrate that these RNAs in the expanded sample are detected with high accuracy and near 100% detection efficiency. In addition, we also demonstrate the ability to perform simultaneous MERFISH RNA imaging and immunofluorescence imaging of proteins in these expanded samples.

## Results

### Effect of RNA density on the detection efficiency of MERFISH measurements

To illustrate the effect of RNA density on multiplexed smFISH measurements, we measured a high-abundance, ~130-RNA library using our previously published 16-bit modified Hamming distance 4 (MHD4) binary code, which allows error detection and correction^7,12,13^. This code includes 140 unique barcodes and we used 129 of them to encode RNAs and kept 11 of them as blank controls that did not correspond to any RNA. Among the 129 targeted RNAs, 106 RNAs were in the abundance range of 40 – 250 copies per cell to increase the total abundance of the library, and the remaining 23 spanned an abundance range of 1 – 1000 copies per cell to quantify performance across different abundances. The total abundance of RNAs in this library is 14-fold higher than the 130-RNA libraries that we have previously measured using the MHD4 code with ~80-90% detection efficiency^7,12,13^.

As in our previous MERFISH measurements^7,12,13^, we labeled the RNAs with two sets of oligonucleotide probes. In the first step, each cellular RNA was hybridized with a complex library of oligonucleotide probes termed encoding probes, which contain targeting sequences that bind cellular RNAs and readout sequences that determine the barcodes of these RNAs. In the second step, the readout sequences, and hence the barcodes, were detected through a series of smFISH measurements, each round with one or more readout probes complementary to one or more readout sequences. We carried out MERFISH measurements in U-2 OS cells using the matrix-imprinting-based clearing method to reduce the fluorescence background as previously published^13^. Briefly, the cells were fixed, permeabilized, labeled with encoding probes for the 129 RNA species as well as acrydite-modified poly-dT LNA probes that target polyadenylated (polyA) RNAs. We then embedded the cells in a non-expandable, polyacrylamide gel and the polyA-containing mRNAs were anchored to the gel through the poly-dT probes. Next, we removed cellular proteins and lipids by Proteinase K digestion and detergent extraction to remove fluorescence background and hence clear the samples. After this clearing, eight rounds of two-color smFISH measurements were performed to read out the 16-bit barcode for each RNA. Because the heights of the cells were greater than the thickness of a single optical section, we imaged the sample with multiple z-sections to ensure that >90% of RNA molecules within the cells were detected.

Because of the high molecular density associated with this RNA library, a substantial fraction of RNA molecules generated spatially overlapping smFISH signals in each round of imaging (Fig. 1A, B). As a result, only a small fraction of the RNA molecules was decodable (Fig. 1C). This observation was in contrast to our previous MERFISH measurements on lower-abundance RNA libraries, where the vast majority of RNAs were spatially separated and decodable^7,12,13^. The average copy number per cell detected for the RNAs in this high-abundance library by MERFISH correlated with the RNA abundance measured by bulk RNA sequencing with a Pearson correlation coefficient (r) of 0.6 (Fig. 1D), which is lower than the ~0.8 correlation coefficient value that we observed previously with lower abundance libraries in this and other cell lines^7,12,13^. To determine the detection efficiency, we measured the abundance of 12 genes in this library individually with smFISH in matrix-imprinted-and-cleared samples. Comparison of the copy numbers determined for these genes with MERFISH and smFISH showed that MERFISH only detected 21% ± 4% (average ± s.e.m.) or 16% (median) of the RNA copies determined by smFISH (Fig. 1E), which also contrasts with the ~80-90% detection efficiency that we observed previously for lower abundance RNA libraries^7,12,13^. Thus, as expected, we find that the ability of MERFISH to identify and quantify RNAs drops when the abundance of the measured RNAs lead to substantial overlap in the signals from neighboring RNAs.

**Figure 1.**
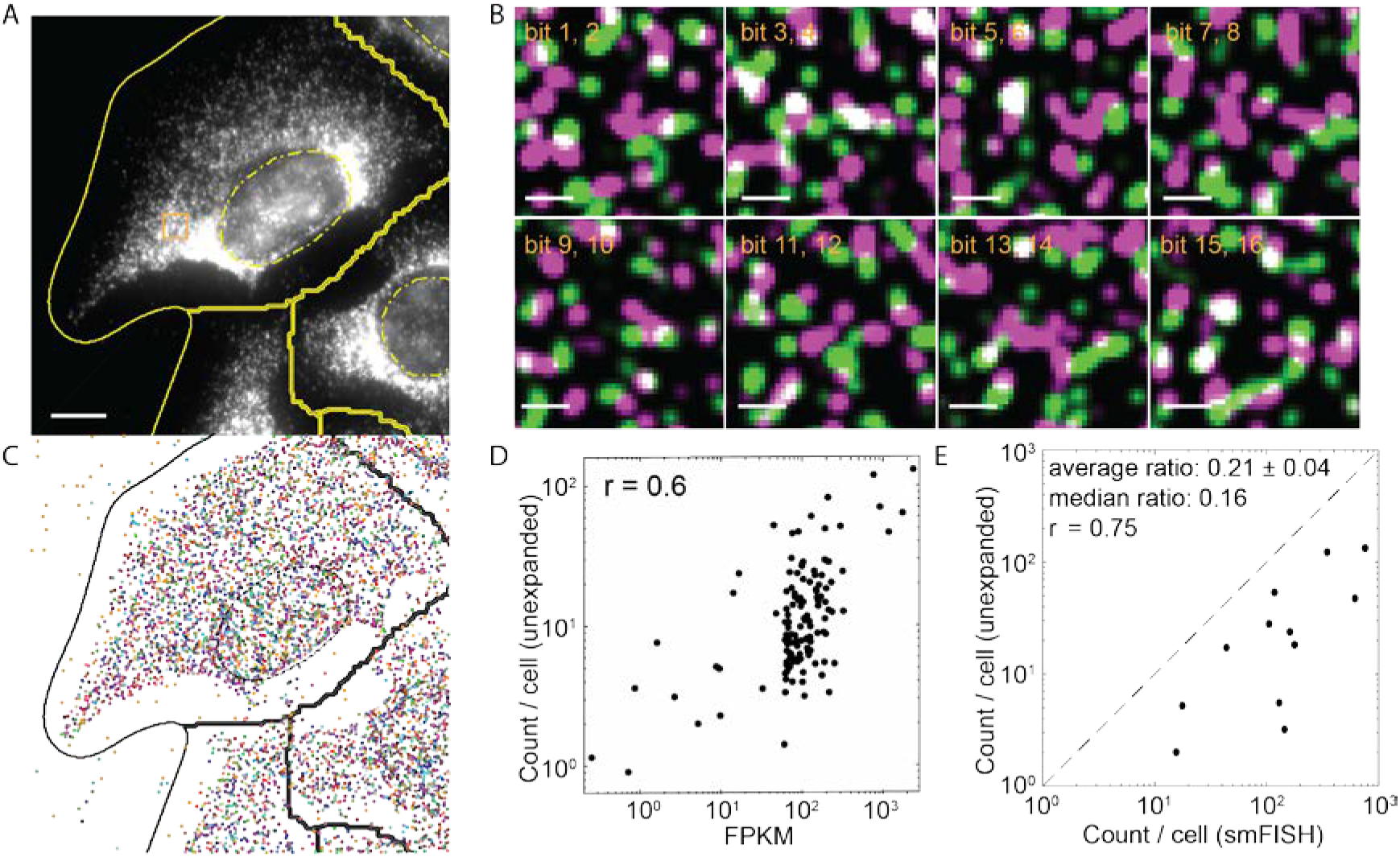
MERFISH measurements of a high-abundance RNA library in unexpanded U-2 OS samples. (**A**) Image of gel-embeded and cleared U-2 OS sample stained with encoding probes for 129 RNAs and a Cy5-labeled readout probe that detects one of the bits of the RNA barcodes at one focal plane of the z-scan. The yellow solid line marks the lines separating neighboring cells, which do not necessarily mark the physical edges of the cells. The yellow dashed line marks the DAPI-stained region of a nucleus. (**B**) Fluorescence images of all 8 rounds of two-color readout imaging for the orange boxed region in (A). Images were deconvolved and Gaussian filtered. Magenta and green represent the Cy5 channel and Alexa 750 channel, respectively. (**C**) The localizations of all decoded RNAs in (A) colored according to their measured binary barcodes. Decoded RNAs across all z-sections are displayed. The black solid line marks the lines separating neighboring cells and the black dashed line marks the DAPI-stained region of a nucleus. (**D**) The average RNA copy numbers per cell for the 129 RNA species determined by MERFISH vs. the abundances as determined by RNA-seq. The Pearson correlation coefficient (r) between the log10 values of MERFISH-determined copy number per cell and RNA-seq determined FPKM value is 0.6. (**E**) The average RNA copy numbers per cell determined by MERFISH vs. those by smFISH for 12 of the 129 RNAs. The Pearson correlation coefficient (r) between the log10 values of MERFISH-determined and smFISH-determined copy numbers is 0.75. The average ratio of the copy number values determined by MERFISH to that determined by smFISH is 0.21 ± 0.04 (s.e.m., *n* = 12 RNA species) and the median ratio is 0.16. The scale bar in (A) represents 10 μm; the scale bars in (B) represent 1 μm.

### High RNA-density MERFISH measurements with expansion microscopy

We reasoned that the reduced MERFISH detection efficiency due to overlapping single-molecule signals could be alleviated by sample expansion, which will increase the distance between molecules and reduce overlap. To test this idea, we fixed and permeabilized the cells, and labeled them with the encoding probes and poly-dT anchoring probes as described above. Then we embedded the cells in an expandable polyelectrolyte gel using a protocol modified from the published ExM methods^20,21^. Afterwards, cells were digested by Proteinase K and treated with detergent to remove proteins and lipids, clearing the sample and facilitating gel expansion. The gel was then expanded in a low salt buffer and finally embedded again in a non-expandable polyacrylamide gel to stabilize it in the expanded state (Fig. 2A). There are several notable differences between our protocol and the previous ExM protocol used for smFISH^21^. First, in the previous protocol, RNAs were randomly modified with a chemical crosslinker that allowed them to be covalently linked to the expandable gel. To ensure that the majority of RNAs contain at least one anchor, Poisson statistics will dictate that most RNAs will be labeled with more than one anchor. Thus, it is possible that RNAs are connected to the gel at multiple points and stretched during expansion. Here, we anchored each RNA to the gel at a single location—the poly(A) tail. This difference in anchoring geometry could potentially improve separation of the nearby RNA molecules. Second, we found that MERFISH encoding probes did not reliably penetrate the gel after expansion and stabilization, perhaps due to the increased length of these probes compared to those used for conventional smFISH. Thus, we stained the samples with encoding probes before embedding in the gel. By contrast, we found that the short MERFISH readout probes penetrated this gel without a problem, allowing the sequential rounds of smFISH to be performed in expanded samples. Third, to stabilize the already bound encoding probes on RNAs, we utilized a low salt buffer (0.5x saline-sodium citrate (SSC)), instead of water, for the expansion of the gel. Our measurement of the gel volume shows that dialysis in 0.5x SSC produced 2.3-fold expansion in each dimension and 12-fold expansion in volume, which is smaller than that reported previously for expansion in water^20,21^. We note that as long as the expansion is sufficient to separate neighboring RNA molecules, a lower expansion factor has the advantage of allowing faster imaging.

**Figure 2.**
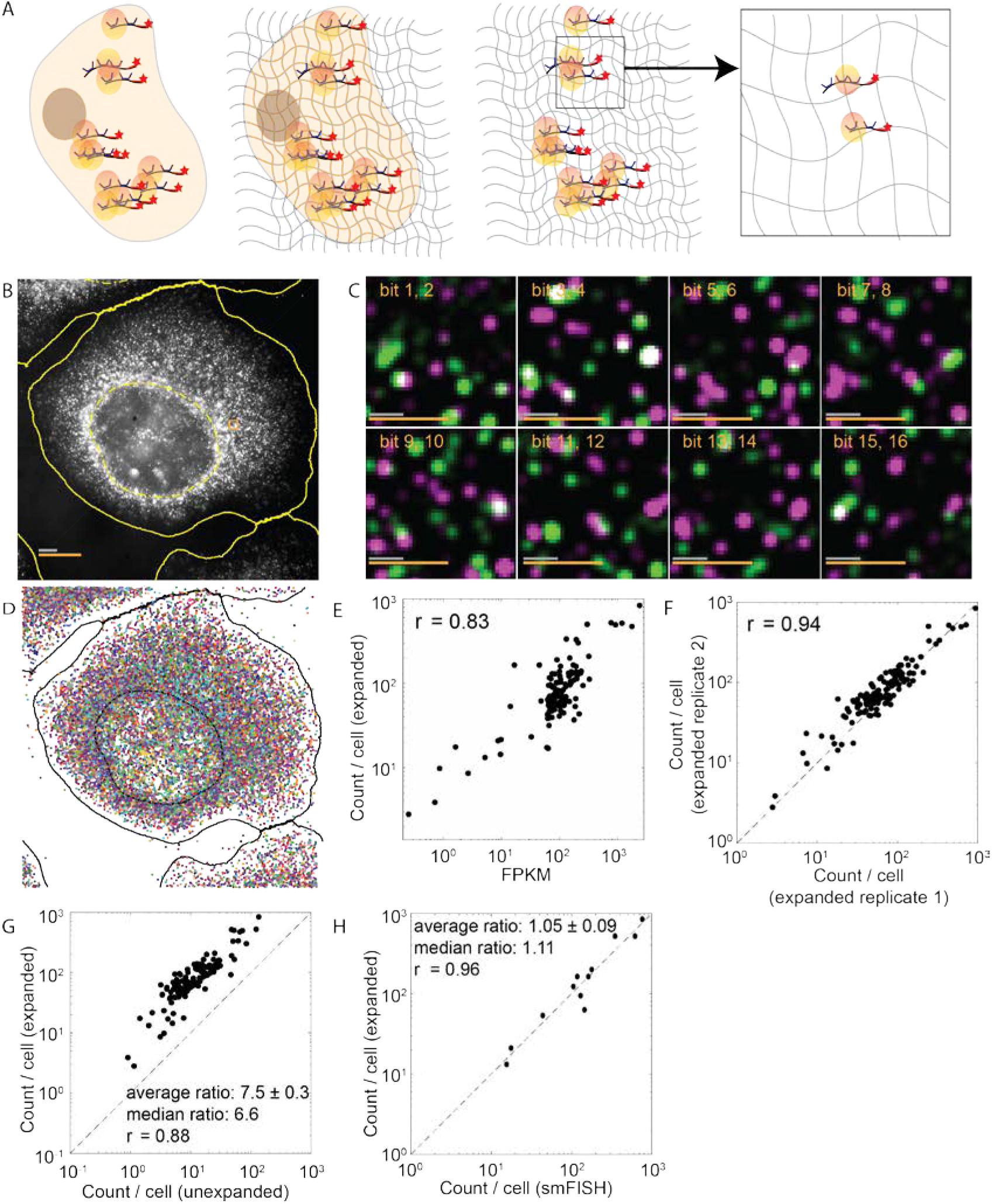
MERFISH measurements of a high-abundance RNA library in expanded U-2 OS cells. (**A**) Schematic representation of the gel embedding and expansion process for MERFISH imaging. (**B**) Image of an expanded U-2 OS sample stained with encoding probes for 129 RNAs and a Cy5-labeled readout probe that detects one of the bits of the RNAs barcodes at one focal plane of the z-scan. (**C**) Fluorescence images of all 8 rounds of two-color readout imaging for the orange boxed region in (B). Magenta and green represent the Cy5 channel and Alexa 750 channel, respectively. (**D**) The localizations of all decoded RNAs in (B) colored according to their measured binary barcodes. Decoded RNAs across all z-sections are displayed. (**E**) The average RNA copy numbers per cell for the 129 RNA species determined by MERFISH vs. the abundances as determined by RNA-seq. The Pearson correlation coefficient (r) based on the log10 values of RNA abundances is 0.83. (**F**) The average RNA copy numbers per cell for the 129 RNA species detected in one expanded sample vs. a replicate sample. The Pearson correlation coefficient (r) is 0.94. (**G**) The average RNA copy numbers per cell for the 129 RNA species determined by expansion MERFISH vs. those by MERFISH in unexpanded samples. The Pearson correlation coefficient (r) is 0.88. The copy numbers per cell detected in expanded samples were 7.5 ± 0.3 fold (average ± s.e.m.) or 6.6 fold (median) of those detected in unexpanded samples for these 129 RNA species. (**H**) The average RNA copy numbers per cell determined by MERFISH on expanded samples vs. those by smFISH measurements on unexpanded samples for 12 of the 129 RNAs. The Pearson correlation coefficient (r) is 0.96. The average ratio between MERFISH and smFISH results is 1.05 ± 0.09 (average ± s.e.m., *n* = 12 RNA species) and the median ratio 1.11. The scale bars in (B) and (C) represent 10 μm and 1 μm, respectively, with the grey scale bars representing the scales in the expanded sample and the orange scale bars representing the scales calculated back to sizes before expansion.

Notably, in the expanded samples, individual RNA molecules became substantially better resolved (Fig. 2B, C) and many more molecules were successfully decoded (Fig. 2D). Compared to the results from unexpanded samples, the average copy number per cell detected for these RNAs by MERFISH correlated better with the RNA abundance measured by RNA-seq with an increased Pearson correlation coefficient of 0.83 (Fig. 2E). The copy number per cell results are highly reproducible between replicates of MERFISH experiments (Fig. 2F).

To quantify the improvement in detection efficiency with expansion, we compared the average MERFISH counts for individual RNA species per cell between expanded and unexpanded samples. We found that the copy numbers per cell for the 129 RNA species detected in expanded samples correlated strongly with those in the unexpanded samples with a high Pearson correlation coefficient of 0.88 (Fig. 2G). However, the copy numbers per cell detected in expanded samples were 7.5 ± 0.3 fold (average ± s.e.m.) or 6.6 fold (median) higher than those detected in unexpanded samples for these 129 RNA species, suggesting that the detection efficiency for the expanded sample is substantially higher compared to the unexpanded sample for this high-abundance library. Indeed, comparison with smFISH measurements shows that the copy numbers per cell determined by MERFISH in the expanded samples is 105% ± 9% (average ± s.e.m.) or 111% (median) of those determined by smFISH (Fig. 2H), indicating that the detection efficiency after expansion is close to 100%. Compared to the 16% median value described earlier for the unexpanded samples, the 111% median value obtained here for the expanded samples suggests a 6.9-fold increase in the detection efficiency, which is consistent with the 6.6-fold increase in the median copy numbers detected in expanded samples as compared to unexpanded samples. We do not make such comparison for the average values because, for broad distributions, the arithmetic mean values of ratio do not propagate mathematically.

### Incorporating immunofluorescence imaging into MERFISH experiments of expanded samples

Cells are comprised of different structures and compartments, and immunofluorescence is a powerful technique to visualize specific subcellular structures and compartments. We thus tested whether combination of immunofluorescence and MERFISH imaging is possible in the expanded samples. It has been previously demonstrated that ExM can be used to image immunostained samples, using oligo-conjugated antibodies and complementary probes with a methacryloyl group to incorporate immuno-labeling signals into the gel^20^ or via direct crosslinking of protein targets or antibodies to the gel^25–27^. When the oligo-labeling approach is utilized, the positions of the protein targets are detected by hybridization of fluorophore-labeled complementary oligos, which we reason can be incorporated into our MERFISH readout measurements. To demonstrate this ability, we performed immunostaining of the protein targets with primary antibody and oligo-labeled secondary antibody after hybridization of the cells with the MERFISH encoding probes and poly-dT anchor probes, and then embedded the labeled sample in an expandable gel. We added acrydite modification to the oligo on the antibodies so that it can be incorporated into the polymer gel during the embedding step. After digestion, gel expansion and second embedding in a non-expandable gel as described in the previous section, we performed the MERFISH readout procedure to first detect the readout sequences in the RNA encoding probes and then with an additional round of FISH detection to read out the oligo sequence representing the protein target.

For demonstration purpose, we immunostained cadherin in cultured U-2 OS cells, which were also stained for the same high-abundance RNA library described in the earlier sections. We observed both specific staining of cadherin on the cell periphery (Fig. 3A) as well as clearly resolved smFISH spots during each readout round in the same cells (Fig. 3B, C). The combination with immunofluorescence did not affect MERFISH imaging quality. We observed the same high correlation of MERFISH results and the RNA-seq results with a Pearson correlation coefficient of 0.83 (Fig. 3D). The average copy numbers per RNA species per cell detected in immunostained samples correlated strongly with those in detected samples not subjected to immunostaining, with a Pearson correlation coefficient of 0.95 (Fig. 3E). On average, the ratio of copy numbers of individual RNA species per cell between immunostained and non-immunostained samples is 0.99 ± 0.03 (average ± s.e.m.), indicating negligible impairment in performance of MERFISH when combined with immunofluorescence. The staining of cadherin in expanded and MERFISH labeled samples also looked similar to cadherin staining in control U-2 OS samples immuostained directly after fixation without any gel-embedding, expansion or MERFISH RNA labeling (Fig. 3F). Quantitatively, we compared the cadherin intensity per unit length on the cell periphery (normalized to the actual length before expansion) in expanded MERFISH samples versus the control samples not subjected to any MERFISH RNA labeling and gel embedding, and obtained very similar results (Fig. 3G).

**Figure 3.**
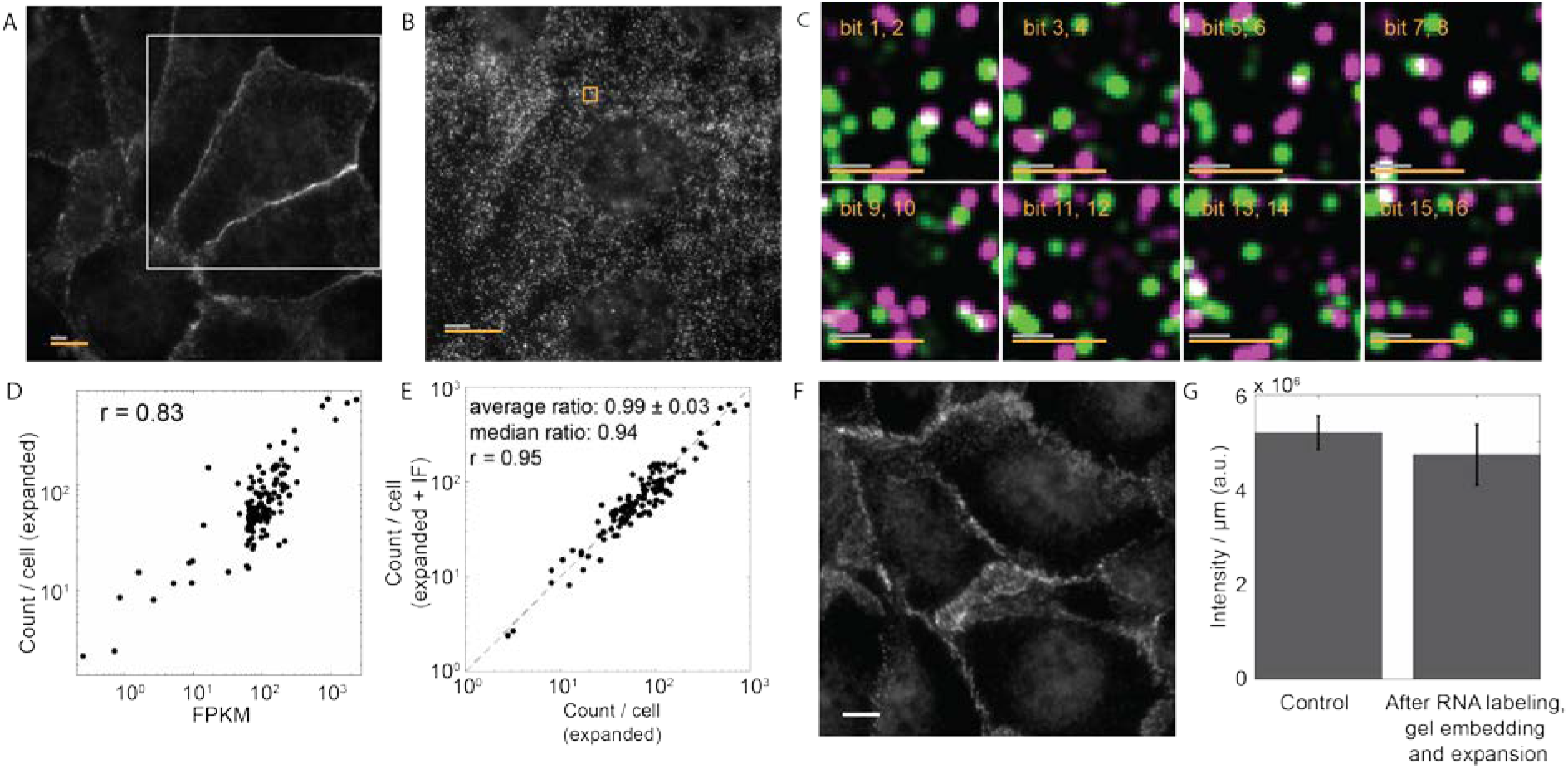
Combining immunofluorescence imaging and MERFISH RNA imaging in expanded samples. (**A**) Image of an expanded U-2 OS sample stained with MERFISH encoding probes for 129 RNAs, cadherin primary antibodies, oligo-conjugated secondary antibodies and visualized with a Cy5-conjugated complementary probe that detects the secondary antibodies. (**B**) Image of the same expanded sample visualized with a Cy5-labeled readout probe that detect one of the bits of the RNA barcodes at one focal plane for the white boxed region in (A). (**C**) Fluorescence images of all 8 rounds of two-color readout imaging for the orange boxed region in (B). Magenta and green represent the Cy5 channel and Alexa 750 channel, respectively. (**D**) The average RNA copy numbers per cell for the 129 RNA species determined by MERFISH vs. the abundances as determined by RNA-seq. The Pearson correlation coefficient (r) based on the log10 values of RNA abundances is 0.83. (**E**) The average RNA copy numbers per cell for the 129 RNA species determined by MERFISH in immunostained samples vs. those in samples that are not immunostained. The Pearson correlation coefficient (r) is 0.95. The average ratio between the RNA copy numbers determined in the two experiments is 0.99 ± 0.03 (average ± s.e.m., *n* = 129 RNA species) and the median ratio is 0.94. (**F**) Image of unembedded U-2 OS cells stained with the same cadherin primary antibodies and oligo-conjugated secondary antibodies as well as a Cy5-conjugated complementary probe right after fixation and permeabilization without any MERFISH RNA labeling and gel embedding. (H) Cadherin intensity per unit length of cell periphery (normalized to the actual length before expansion) in expanded MERFISH samples and in control immunostained samples that have not been subject to any MERFISH RNA labeling or gel embedding. Error bars are s.e.m. (*n* = 20). The scale bars in (A) and (B) represent 10 μm and the scale bars in (C) represent 1 μm, with the grey scale bars representing the scales in the expanded sample and the orange scale bars representing the scales calculated back to sizes before expansion. The white scale bar in (F) represents 10 μm.

## Discussion

In this paper, we have demonstrated an approach to combine MERFISH and ExM to measure high-abundance RNA libraries. We showed that when the total RNA density is high, the overlap between signals from nearby RNA molecules reduced the MERFISH detection efficiency and that sample expansion can overcome this overlapping problem and substantially increase the detection efficiency. Specifically, for an RNA library with total RNA abundance that is 14-fold higher than our previously measured libraries, the MERFISH detection efficiency dropped to ~20% in unexpanded samples, and expansion recovered the near 100% detection efficiency of MERFISH.

Each mammalian cell has tens of thousands of different RNA species. We have previously demonstrated MERFISH imaging of up to 1,000 RNA species in single cells^7^. To increase the number of RNA species that can be simultaneously imaged in single cells while maintaining a similar number of imaging rounds, the density of RNAs imaged per round will increase and will likely become an important limiting factor in the number of RNAs that can be imaged. We anticipate that this expansion MERFISH approach should facilitate a substantial increase in the number of RNA species that can be measured in single cells.

We also demonstrated the combination of MERFISH with immunofluorescence in these expanded samples, which can provide important information on the cellular context for transcriptome analysis. In conventional immunofluorescence with fluorophore-conjugated antibodies, the number of spectrally-distinct fluorophores limits the number of targets that can be studied simultaneously. The use of oligo-conjugated antibodies potentially allows visualization of many proteins in sequential rounds of hybridization and imaging using just one or a few spectrally distinct fluorophores^28–31^. We thus envision such combination of MERFISH and immunofluorescence imaging may allow a combined transcriptomic and proteomic measurements simultaneously in single cells.

## Materials and Methods

### Design of the encoding probes

The MERFISH encoding probes were designed using the same 16-bit Hamming-weight-4 Hamming-distance-4 code with 140 possible barcodes as previously published^7,12^. In this encoding scheme, all barcodes used are separated by a Hamming distance of at least 4, and hence at least four bits must be read incorrectly to change one valid barcode to another. A constant Hamming weight (i.e. the number of “1” bits in each barcodes) of 4 is used to avoid potential measurement bias due to differential rates of “1” to “0” and “0” to “1” errors^7,12^. The encoding probe set that we used contained 92 encoding probes per RNA. Each encoding probe was comprised of a 30-nt target region designed using a pipeline published previously^12^, flanked by two 20-nt readout sequences randomly selected out of the four ones assigned to each RNA, one 20-nt priming region at the 5’ end and another 20-nt priming region using the reverse complement of the T7 promoter at the 3’ end. Additional adenosine nucleotide spacers were added between readout sequences and target regions to prevent target regions from combining with Gs from adjacent sequences to form G quadruplets. The priming region at the 5’ end had a thymine at the end, which was put at the junction of the priming regions at the 5’ end and the encoding region. This was designed to incorporate a uracil on the forward primer used in the reverse transcription step of probe construction, which can be cleaved by Uracil-Specific Excision Reagent (USER) Enzyme^32^. The cleavable primer design together with using the reverse complement of the T7 promoter as the second primer allowed us to remove these priming regions from the final probes, producing encoding probes with a length of 72 nt compared to 112 nt using previous approaches^7,12,13^. The reduction in probe length may facilitate penetration of probes and reduce non-specific binding. To include some abundant but relatively short RNAs in our library, instead of designing probes targeting distinct regions of RNAs, we allowed probes to share up to 20 nt with another probe. 92 target regions per RNA were selected randomly out of all potential target regions of an RNA. By allowing up to a 20-nt overlap between neighboring encoding probes, this design allowed RNAs as short as 1,200 nt to be targeted by 92 encoding probes with a 30-nt targeting sequence. Because a given cellular RNA is typically bound by less than one third of the 92 encoding probes, we reason that the encoding probes with overlapping targeting regions would not substantially interfere with each other but would partially compensate for reduced binding due to local inaccessible regions on the target RNA (e.g. secondary structure) or loss of probe during synthesis. The same 20-nt, three-letter readout sequences were used as previously published^12^.

### Construction of the encoding probes

The encoding probe set was amplified from complex oligonucleotide pools. Briefly, we amplified the oligopools (CustomArray) via limited-cycle PCR to make in vitro transcription templates, converted these templates into RNA via in vitro transcription (New England Biolabs), and converted the RNA back to DNA via reverse transcription (Maxima RT H-, Thermo Fisher Scientific) as previously published^12^. The probes were then digested by USER Enzyme^32^ (New England Biolabs) at a dilution of 1:30 (vol/vol) incubated at 37 °C for 24h to cleave off the priming region at the site of a uracil between a priming region at the 5’ end and the target region. After that, DNA was purified via alkaline hydrolysis to remove RNA and column purification (Zymo Research). The final probes were resuspended in RNAase-free water and stored at −20 °C.

### Construction of the readout probes

The three-letter readout sequences were designed as published previously^12,33^. Readout probes conjugated to the desired dye via a disulfide linkage were synthesized and purified by Bio-synthesis, Inc., resuspended immediately in Tris-EDTA (TE) buffer, pH 8 (Thermo Fisher) to a concentration of 100 μM and stored at −20 °C. To reduce the number of freeze-thaw cycles, 1-μM aliquots were made in TE buffer and stored at −20 °C.

### Oligo conjugation to secondary antibodies

Oligonucleotides containing the complementary sequence of the desired readout probe were conjugated to secondary antibodies via a combination of NHS-ester and copper-free click chemistries similar to a published method^34^. First, secondary antibodies were labeled with a copper-free click crosslinking agent using NHS-ester chemistry. Specifically, azide preservative was removed from the unconjugated Donkey Anti-Rabbit secondary antibodies (Thermo Fisher Scientific) using a spin-column based dialysis membrane (Amicon, 100kDa molecular weight cut off) according to the manufacturer’s instructions. DBCO-PEG5-NHS Ester (Kerafast) was diluted to a concentration of 10 μM in anhydrous dimethyl sulfoxide (DMSO) (Thermo Fisher Scientific). 2 μL of the solution was then combined with 100 μL of 2 mg/mL of the antibody in 1× phosphate-buffered saline (PBS). This reaction was incubated at room temperature for 1 hour and then terminated via a second round of purification using the Amicon columns as described above. The average number of DBCO crosslinkers per antibody was determined via the relative absorption of the sample at 280 nm (antibody) and 309 nm (DBCO). On average the procedure described above produced ~7 DBCO per antibody.

Oligonucleotide probes containing the desired sequence as well as a 5’-acrydite, to allow cross-linking to the polymer gel, and a 3’-azide, to allow cross-linking to the DBCO-labeled antibodies, were ordered from IDT and suspended to 100 μM in 1×PBS. 20 μL of the appropriate oligonucleotide was then added to 100 μL of the DBCO-labeled antibodies at a final concentration of ~2 mg/mL. This reaction was incubated at 4C for at least 12 hours. Labeled antibodies were not further purified as residual oligonucleotides, not conjugated to antibodies, were readily washed away from samples.

### Cell culture and fixation

U-2 OS cells (ATCC) were cultured with Eagle’s Minimum Essential Medium (ATCC) containing 10% (vol/vol) fetal bovine serum (FBS) (Thermo Fisher Scientific). Cells were plated on 40-mm-diameter, no.1.5 coverslips (Bioptechs) at 350,000 cells per coverslip and were incubated in Petri dishes at 37 °C with 5% CO_2_ for 48 h. Cells were fixed for 15 min in 4% (vol/vol) paraformaldehyde (PFA) (Electron Microscopy Sciences) in 1× PBS at room temperature, washed three times with 1× PBS, permeabilized for 10 min with 0.5% (vol/vol) Triton X-100 (Sigma) in 1× PBS at room temperature, and washed once with 1× PBS.

### Encoding probe staining

Permeabilized cells were incubated for 5 min in encoding wash buffer comprising 2× saline-sodium citrate (SSC) (Ambion) and 30% (vol/vol) formamide (Ambion). Then 30 μL of ~300 μM encoding probes and 3.3 μM of poly-dT LNA anchor probe (a 20-nt sequence of alternating dT and thymidine-locked nucleic acid (dT+) with a 20-nt reverse complement of a readout sequence and a 5’-acrydite modification (Integrated DNA Technologies)) in encoding hybridization buffer was added to the surface of Parafilm (Bemis) and was covered with a cell-containing coverslip. Encoding hybridization buffer was composed of encoding wash buffer supplemented with 0.1% (wt/vol) yeast tRNA (Life Technologies), 1% (vol/vol) murine RNase inhibitor (New England Biolabs), and 10% (wt/vol) dextran sulfate (Sigma). Samples were incubated in a humid chamber inside a hybridization oven at 37 °C for 40 h. Cells then were washed with encoding wash buffer and incubated at 47 °C for 30 min; this washing step was repeated once.

### Immunostaining

For immunofluorescence only, cells were fixed and permeabilized as described in the “Cell culture and fixation” Section. Samples were first blocked at room temperature for 30 min in blocking buffer consisted of 4% (wt/vol) UltraPure BSA (Thermo Fisher Scientific) in 2× SSC supplemented with 3% (vol/vol) RNasin Ribonuclease inhibitor (Promega), 6% (vol/vol) murine RNAase inhibitor and 1 mg/ml yeast tRNA. Samples were then incubated with primary antibodies (anti-pan Cadherin, Abcam) in blocking buffer at a concentration of 2 μg/ml for 1h at room temperature, and washed three times with 2× SSC for 10 min each. Samples were then incubated with oligo-labeled secondary antibodies in blocking buffer at a concentration of 3.75 μg/ml for 1h at room temperature, then washed with 2× SSC three times for 10min each. Samples were fixed again with 4% (vol/vol) PFA in 2× SSC for 10 min and washed three times with 2× SSC.

To combine MERFISH with immunofluorescence, samples were first stained with encoding probes and washed as described in the “Encoding probe staining” section above. To stabilize these probes, samples were then briefly post-fixed with 4% (vol/vol) PFA in 2× SSC at room temperature for 10 min and washed three times with 2× SSC. Samples were then stained for immunofluorescence as described above.

### Gel embedding, digestion and clearing for unexpanded samples

Stained samples on silanized coverslips treated as in “Silanization of coverslips for unexpanded samples” Section (see below), were first incubated for 5 min with a de-gassed polyacrylamide solution, consisted of 4% (vol/vol) of 19:1 acrylamide/bis-acrylamide (BioRad), 60 mM Tris·HCl pH 8 (Thermo Fisher Scientific), 0.3 M NaCl (Thermo Fisher Scientific), 0.2% (vol/vol) Tetramethylethylenediamine (TEMED) (Sigma) and a 1:25,000 dilution of 0.1-μm-diameter carboxylate-modified orange fluorescent beads (2% solids, Life Technologies). The beads served as fiducial markers for the alignment of images taken across multiple rounds of smFISH imaging. The polyacrylamide solution was kept on ice and further supplemented with ammonium persulfate (Sigma) at a final concentration of 0.2% (wt/vol).

50 μL of this gel solution was added to the surface of a glass plate (TED Pella) that had been pretreated for 5 min with 1 mL GelSlick (Lonza) so as not to stick to the polymer gel. Samples were aspirated, dried quickly with KimWipes (Kimtech) from the edge of the coverslips, and gently inverted onto this 50-μL droplet to form a thin layer of solution between the coverslip and the glass plate. The solution was then allowed to polymerize for 2 h at room temperature in a home-built chamber filled with nitrogen. The coverslip and the glass plate were then gently separated, and the PA film was washed once with a digestion buffer consisted of 2% (wt/vol) Sodium dodecyl sulfate (SDS) (Thermo Fisher Scientific), 0.5% (vol/vol) Triton X-100 in 2× SSC. Our digestion buffer is different from published previously^13,20,21^. We used 2% (wt/vol) SDS to facilitate lipid removal. After the wash, the gel was covered with digestion buffer supplemented with 1% (vol/vol) Proteinase K (New England Biolabs). The sample was digested in this buffer for >12 h in a humidified, 37 °C incubator and then washed three times with 2× SSC for 15 min each on a rocker. MERFISH measurements were either performed immediately or the sample was stored in 2× SSC supplemented with 0.1% (vol/vol) murine RNase inhibitor at 4 °C for no longer than 48 h.

### Silanization of coverslips for unexpanded samples

To stabilize the polymer film, coverslips were silanized as published previously^13,35^. Briefly, 40-mm-diameter #1.5 coverslips (Bioptechs) were washed for 30 min via immersion in a 1:1 mixture of 37% (vol/vol) hydrochloric acid (Sigma) and methanol (Sigma) at room temperature. Coverslips were then rinsed three times in deionized water and once in 70% (vol/vol) ethanol. Coverslips were blown dry with nitrogen gas and then immersed in 0.1% (vol/vol) triethylamine (Millipore) and 0.2% (vol/vol) allyltrichlorosilane (Sigma) in chloroform for 30 min at room temperature. Coverslips were washed once each with chloroform and ethanol and then blow dry with nitrogen gas. Silanized coverslips were then be stored at room temperature in a desiccated chamber overnight before use to dehydrate the silane layer.

### Gel embedding, digestion and clearing, and expansion for expanded samples

The embedding and expansion protocol was modified based on previously published methods^20,21^. Monomer solution consisted of 2 M NaCl, 7.7% (wt/wt) sodium acrylate (Sigma), 4% (vol/vol) of 19:1 acrylamide/bis-acrylamide and 60 mM Tris-HCl pH 8 was prepared and frozen in aliquots at −20 °C. Monomer solution was thawed, degassed and cooled to 4 °C on ice before use. TEMED with a final concentration of 0.2% (vol/vol) and a 1:5,000 dilution of 0.1-μm-diameter carboxylate-modified orange fluorescent beads was added to the solution. The beads served as fiducial markers for the alignment of images taken across multiple rounds of smFISH imaging. Stained samples were incubated in the solution for 5 min at room temperature. The solution was then kept on ice and further supplemented with ammonium persulfate at a final concentration of 0.2% (wt/vol).

The casting of a thin polymer film and polymerization was performed the same as described in the “Gel embedding, digestion and clearing for unexpanded samples” Section. After polymerization, the coverslip and the glass plate were gently separated. The gel film on the coverslip was washed once with the digestion buffer and trimmed to desired sizes using a razor blade. Digestion was performed the same as described in the “Gel embedding, digestion and clearing for unexpanded samples” Section. The gel would expand ~1.5 fold during digestion.

After digestion, samples were expanded in 0.5× SSC buffer supplemented with 0.2% (vol/vol) Proteinase K at room temperature. Proteinase K was added to maintain samples in an RNAase free environment and to digest away any newly exposed proteins. The buffer was changed every 30 min until samples no longer expanded (typically ~2h). Expanded gels were re-embedded in polyacrylamide gel to stabilize the gel for sequential rounds of readout probe hybridization and imaging. Briefly, samples were incubated in re-embedding solution composed of 4% (vol/vol) of 19:1 acrylamide/bis-acrylamide with 30 mM NaCl, 6 mM Tris ·HCl pH 8 and 0.2% (vol/vol) of TEMED for 20 min at room temperature on a rocker. The re-embedding solution was then kept on ice and further supplemented with ammonium persulfate at a final concentration of 0.2% (wt/vol). Gels were placed on a bind-silane-treated coverslip prepared using the protocol described in “Bind-silane treatment of coverslips” section (see below), rinsed with the solution and dried quickly with KimWipes. Coverslips with gels were put in a home-built nitrogen chamber, covered a glass plate and allowed to polymerize at room temperature for 1h. The salt concentrations in the buffers utilized for expansion and re-embedding were determined so that the encoding probes are well maintained on the RNA during these processes. The salt concentration may need to be adjusted when different probe sets are used.

### Bind-silane treatment of coverslips

40-mm-diameter, no.1.5 coverslips (Bioptechs) were sonicated in 1M potassium hydroxide for 30 min, wash three times with deionized water and sonicated again in 70% (vol/vol) ethanol for 30min. Coverslips were silanized using a modified version of published protocols^21,36^. Briefly, coverslips were immersed in a solution composed of 5% (vol/vol) glacial acetic acid (Sigma) and 0.38% (vol/vol) bind-silane (GE Healthcare) in 99% (vol/vol) ethanol for 1h at room temperature. After being quickly washed with 70% (vol/vol) ethanol three times, coverslips were put into a 60 °C oven until dried completely. Coverslips can be stored in seal containers with desiccants for up to a month.

### Sequential rounds of readout probe staining and imaging

To facilitate choosing the right focal plane for imaging, embedded samples were hybridized in dish with the first pair of two-color readout probes in hybridization buffer composed of 2× SSC, 5% (vol/vol) ethylene carbonate (Sigma), 0.1% (vol/vol) murine RNase inhibitor in nuclease-free water, and 3 nM of the appropriate readout probes, for 30min (expanded samples) or 10min (unexpanded samples) at room temperature and washed for 20min (expanded samples) or 7 min (unexpanded samples) in wash buffer composed of 2× SSC and 10% (vol/vol) ethylene carbonate in nuclease-free water. Samples were then washed with 2× SSC once, stained with 4’,6-Diamidino-2-Phenylindole, Dihydrochloride (DAPI) (Thermo Fisher Scientific) at 10μg/ml in 2× SSC for 10min, and washed 3 times in 2× SSC for 5 min each.

Samples were then mounted into a flow chamber and buffer exchange through this chamber was controlled via a home-built fluidics system composed of three computer-controlled eight-way valves (Hamilton) and a computer-controlled peristaltic pump (Gilson) as published previously^7,12^. Flow and incubation time was increased for expanded samples (besides the hybridization and wash time as described in the previous section, tris(2-carboxyethyl)phosphine hydrochloride (TCEP) (Sigma) incubation time was increased to 30 min and all buffer exchange time to 7 min) to allow diffusion to reach equilibrium inside of the gel.

In order to accurately compare the counts per cell between different samples, we performed z-scanning to image more than 90% of target RNAs in a cell. Specifically, we performed optical sectioning at discrete imaging planes across the cell with a step size of 0.5 μm and quantified distribution of smFISH signals as a function of z position. We found >90% of RNAs located in the first 5 μm-depth volume in unexpanded samples and in the first 12 μm-depth volume in expanded samples from the surface of coverslips. Thus, we performed z-stack imaging by scanning a 5 μm-depth volume for unexpanded samples and a 12 μm-depth volume for expanded samples with a step size of 1 μm. The scanning in z direction was controlled by a Nano-F200 nanopositioner (Mad City Labs).

Sequential MERFISH imaging and signal removal was carried out on a high-throughput imaging platform as published previously^12^. Briefly, after hybridization, samples were imaged with a FOV area of 223 × 223 μm utilizing a 2,048 × 2,048 pixel, scientific complementary metal-oxide semiconductor (sCMOS) camera (Zyla 4.2; Andor) in combination with a high numerical aperture (NA = 1.3) and a high-magnification (60×) silicone oil objective (UPLSAPO 60×S2; Olympus). After imaging ~100-400 FOVs, the fluorescence of the readout probes was extinguished by incubating the sample in a reductive cleavage buffer composed of 2× SSC and 50 mM TCEP. The hybridization, imaging and chemical cleavage process was repeated eight times for MERFISH-only samples and nine times for MERFISH+IF samples, with 405-nm DAPI channel imaged in conjunction with the first round of readout imaging.

### Image registration and decoding

Registration of images of the same FOV across imaging rounds as well as decoding of the RNA barcodes was conducted using a previously published analysis pipeline^12^. Briefly, z-stacks at each location from different imaging rounds were corrected for lateral offsets based on the location of fiducial beads. Each focal plane of the corrected z-stacks were high-pass filtered to remove background, deconvolved to tighten RNA spots, and then Gaussian-pass filtered to facilitate connecting signals from one imaging round to another. To correct for differences in the brightness between color channels, images were first normalized by equalizing their intensity histograms and refined further via iterative decoding trials to remove substantial variation in the fluorescence intensity between different bits. The set of 16 normalized intensity values (corresponding to 16 readout imaging) observed for each pixel in each FOV at each focal plane represented a vector in a 16-dimensional space. The pixel vector was normalized and compared to each of the 140 barcodes in the 16-bit MHD4 code. A pixel was assigned to a barcode if the Euclidean distance between the vector and a barcode was smaller than a given threshold defined by the distance of a single-bit error. Adjacent pixels in all focal planes of the z-stacks were combined into a single putative RNA using a 3-D connectivity array with maximal neighborhood connectivity.

Cells were segmented using a previously published approach^12^. Briefly, the lines separating cells were calculated using the watershed algorithm based on the inverted barcode density with DAPI stained regions as initial seeds.

Computations were split between the Odyssey cluster supported by the FAS Division of Science, Research Computing Group at Harvard University and a desktop server that contained two 10-core Intel Xeon E5-2680 2.8GHz CPUs and 256 GB of RAM.

### Single-molecule FISH

Each smFISH probe contains a 30-nt target region, designed using a pipeline published previously^12^, and a 20-nt custom-designed readout sequence^12^. We designed 48 probes for each gene. The probes were synthesized from IDT, resuspended at 100 μM per probe in TE buffer, and pooled together for each gene. After permeabilization, cells on silanized coverslips treated as in “Silanization of coverslips for unexpanded samples” Section were incubated for 5 min in encoding wash buffer comprising 2× SSC and 30% (vol/vol) formamide. Then 30 μL of 2 μM smFISH probes and 3.3 μM of poly-dT LNA anchor probe in encoding hybridization buffer was added to the surface of Parafilm and was covered with a cell-containing coverslip. Samples then were incubated in a humid chamber inside a hybridization oven at 37 °C for 24 h. Encoding hybridization buffer was the same as the one used for MERFISH. Cells then were washed with encoding wash buffer and were incubated at 47 °C for 30 min; this washing step was repeated once. Cells were then embedded and cleared using the matrix-imprinting-based clearing method described in “Gel embedding, digestion and clearing for unexpanded samples” Section. Embedded samples were hybridized in dish with a Cy5 20-nt readout probe in hybridization buffer, washed and stained with DAPI as described in “Sequential rounds of readout probe staining and imaging” Section. Images were collected scanning of a 5 μm-depth volume at a step size of 1 μm and 100 FOVs were collected for each smFISH probe set. smFISH spots were detected using the multi-emitter fitting routine 3D-DAOSTORM^37^ for each z section and spots appearing in adjacent z sections at the same x,y location were combined into one.

## Data availability

The datasets that support the finding of this paper are available from the corresponding author upon request.

## Code availability

The software used to analyze the datasets are available from the corresponding author upon request.

## Acknowledgement

We thank Edward S. Boyden (Massachusetts Institute of Technology) for helpful discussions regarding expansion microscopy.

## Author Contributions

G.W., J.R.M., and X.Z. designed research; G.W. performed experiments; G.W and J.R.M. analyzed data; and G.W., J.R.M., and X.Z. wrote the paper.

